# Potent programmable antiviral against dengue virus in primary human cells by Cas13b RNP with short spacer and delivery by virus-like particle

**DOI:** 10.1101/2020.05.17.100388

**Authors:** Ekapot Singsuksawat, Suppachoke Onnome, Pratsaneeyaporn Posiri, Amporn Suphatrakul, Nittaya Srisuk, Rapirat Nantachokchawapan, Hansa Praneechit, Chutimon Sae-kow, Pala Chidpratum, Suradej Hongeng, Panisadee Avirutnan, Thaneeya Duangjinda, Bunpote Siridechadilok

## Abstract

With sequencing as a standard frontline protocol to identify emerging viruses such zika virus and SARS-CoV2, direct utilization of sequence data to program antivirals against the viruses could accelerate drug development to treat their infections. CRISPR-Cas effectors are promising candidates that could be programmed to inactivate viral genetic material based on sequence data but several challenges such as delivery and design of effective crRNA need to be addressed to realize practical use. Here, we showed that virus-like particle (VLP) could deliver PspCas13b-crRNA ribonucleoprotein (RNP) in nanomolar range to efficiently suppress dengue virus infection in primary human target cells. Shortening spacer length could significantly enhance RNA-targeting efficiency of PspCas13b in mammalian cells compared to the natural length of 30 nucleotides without compromising multiplex targeting by a crRNA array. Our results demonstrate the potentials of applying PspCas13b RNP to suppress RNA virus infection, with implications in targeting host RNA as well.

## Introduction

Among the emerging pathogenic human viruses, the majority is RNA viruses. In recent years, large outbreaks of zika viruses, ebola viruses and, currently, SARs-CoV2 have prompted WHO to declare global health emergencies. Effective antivirals and vaccines against these viruses are still being developed. CRISPR-Cas13, bacterial proteins that can be programmed to target specific RNA with crRNA, are promising tools for developing programmable antivirals against RNA viruses (Abbott et al., 2020; Aman et al., 2018a; 2018b; Bawage et al., 2018; Freije et al., 2019). However, several challenges remain to translate it into practical use. Transient delivery of Cas13b-crRNA into target cells is desirable for treating acute viral infection to avoid the toxicity of long-term expression of Cas13 in the cells but efficient delivery Cas13-crRNA into primary cells has not been achieved. Here, we showed that transient delivery of Cas13b RNP in nanomolar range by virus-like particle could efficiently suppress dengue virus infection in several primary human cells. In contrast to previous studies in vitro and in bacteria, we found that shortening spacer length of crRNA in the range of 26-18 nucleotides could enhance knock-down activity by Cas13b in mammalian cells and did not compromise crRNA processing and multiplex targeting capability.

## Results

To characterize the antiviral activity of CRISPR-Cas13 against flavivirus, we first established a stable BHK21 clone (BHK21-Cas13b) that could be induced with doxycycline to express PspCas13b using lentivirus (**Supplementary Figure 1a**). Type I-interferon defective BHK21 cell line was chosen as the host cell for its ability to support high level of dengue virus (DENV) and zika virus (ZIKV) replication(Liu et al., 2006), providing a robust platform to evaluate viral suppression activity. Inducible expression system appeared to provide the stability of Cas13b cassette in BHK21 as we were not able to maintain PspCas13b expression under a constitutive promoter (EF1a) over several cell passaging (data not shown). To test virus suppression, we utilized both fluorescent reporter viruses which provided virus-encoded fluorescent readout of virus replication(Schoggins et al., 2012; Suphatrakul et al., 2018) and natural strains DENV2-16681 and ZIKV-SV0010(Buathong et al., 2015). We found that cytoplasmic PspCas13b with HIV-Rev nuclear export sequence(Cox et al., 2017) could suppress DENV2-mCherry infection with an mCherry-targeting crRNA (mCh3 crRNA, **Supplementary table 1**) and was chosen for subsequent experiments (**Supplementary Figure 1b**). Though the site of flaviviral RNA replication is associated with ER membrane(Neufeldt et al., 2018), localizing PspCas13b to ER with tail-anchor sequences such as SQS or VAMP2(Guna et al., 2018) failed to suppress DENV2-mCherry infection (**Supplementary Figure 1b**). The suppression of DENV2-mCherry by mCh3 crRNA was specific as other reporter DENV2 could not be suppressed by the crRNA in both single-virus infection and co-infection settings (**Supplementary Figures 1c-d**). Using BHK21-Cas13b, we individually tested 51 crRNAs against the targets on DENV2 reporter viruses, DENV2-16681 (NS5 gene), and ZIKV-SV0010 (NS2A gene) (**Supplementary Table 1**). While half of the crRNAs targeting fluorescent reporter genes (6 out of 12) were effective at viral suppression, only a small fraction of tiled crRNAs targeting the viral genes (3 out of 39, **Supplmentary Figure 1e**) could efficiently suppress virus (knock-down ratio < 0.4). We further validated the most efficient DENV2-targeting crRNA, 8681 crRNA, with DENV2-16681. We found that it could reduce virus titer 15 folds relative to the nontarget crRNA at 48 hours post infection (hpi) and was specific to DENV2-16681 as nontarget ZIKV-SV0010 was not affected (**Supplementary Figure 1f**, bar plot). Infected BHK21-Cas13b with 8681 crRNA were still actively dividing at 72 hpi (and potentially maintaining the infectious titer in the media) while most BHK21-Cas13b with mCh3 crRNA were dead from infection with dropping titer (**Supplementary Figure 1f**, bright-field image panels). Thus, PspCas13b with 8681 crRNA could retard DENV2 infection and protect host cell from its cytopathic effect in the absence of functional host antiviral response.

Despite a relatively high and stable expression of CRISPR-Cas13b in our setup (an estimate of 994 ng/100,000 cells when induced with doxycycline at 0.1μg/ml), delivery needs to be improved for practical use. Lentivirus delivery of CRISPR-Cas13 expression cassettes into cells has several limitations. It requires transcription and translation of CRISPR-Cas13 before the effector becomes available to target viral RNA, a process which require several hours and could affect the effectiveness of virus suppression. CRISPR-Cas13 could also be permanently engrafted into the host genome, a process that carries risks of transforming host cells and Cas13-toxicity from prolonged expression. Recently, retrovirus has been re-engineered to deliver protein cargoes and Cas9-gRNA ribonucleoprotein (RNP) into cells in the form of virus-like particle (VLP)(Kaczmarczyk et al., 2011; Mangeot et al., 2019). Cas13 RNP delivery by VLP allows for immediate targeting of virus and avoids genotoxicity. To produce VLP for Cas13b RNP delivery, we fused PspCas13b gene to GAG gene to construct GAG-PspCas13b plasmid and co-transfected it with crRNA, Gag-Pol, and VSV-G plasmids into 293T (**Supplementary Figure 2a**). We first tested VLP delivery of yellow fluorescent protein (YFP) into various dengue natural target cells that include human primary cells such as human dendritic cells (hDC), macrophages, CD14^+^ monocytes(Durbin et al., 2008; Jessie et al., 2004), and hepatocytes(Aye et al., 2014) (iMHC(Pewkliang et al., 2018)). The VLP was effective at delivering YFP to these cells (**Supplementary Figure 1b**). We found that the VSV-G pseudotyped VLP preferentially target CD14^+^ monocytes in human peripheral blood mononuclear cells (PBMC) over B cells and T cells (**Supplementary Figure 1c**). VLP delivery of PspCas13b with 8681 crRNA (**Figure 1a**) could suppress DENV2-16681 infection in human PBMC (**Figure 1b**). The deliveries of ~77ng of Cas13b/100,000 cells (or 1.16 nM) at 2, 6, and 24 hours post infection (hpi) were able to reduce dengue infection by half over the infection course of 48 hours (**Figure 1b**). Together, these results show that VLP can be an effective way to deliver antiviral Cas13b RNP into primary human target cells to reduce dengue infection.

**Figure 1:**
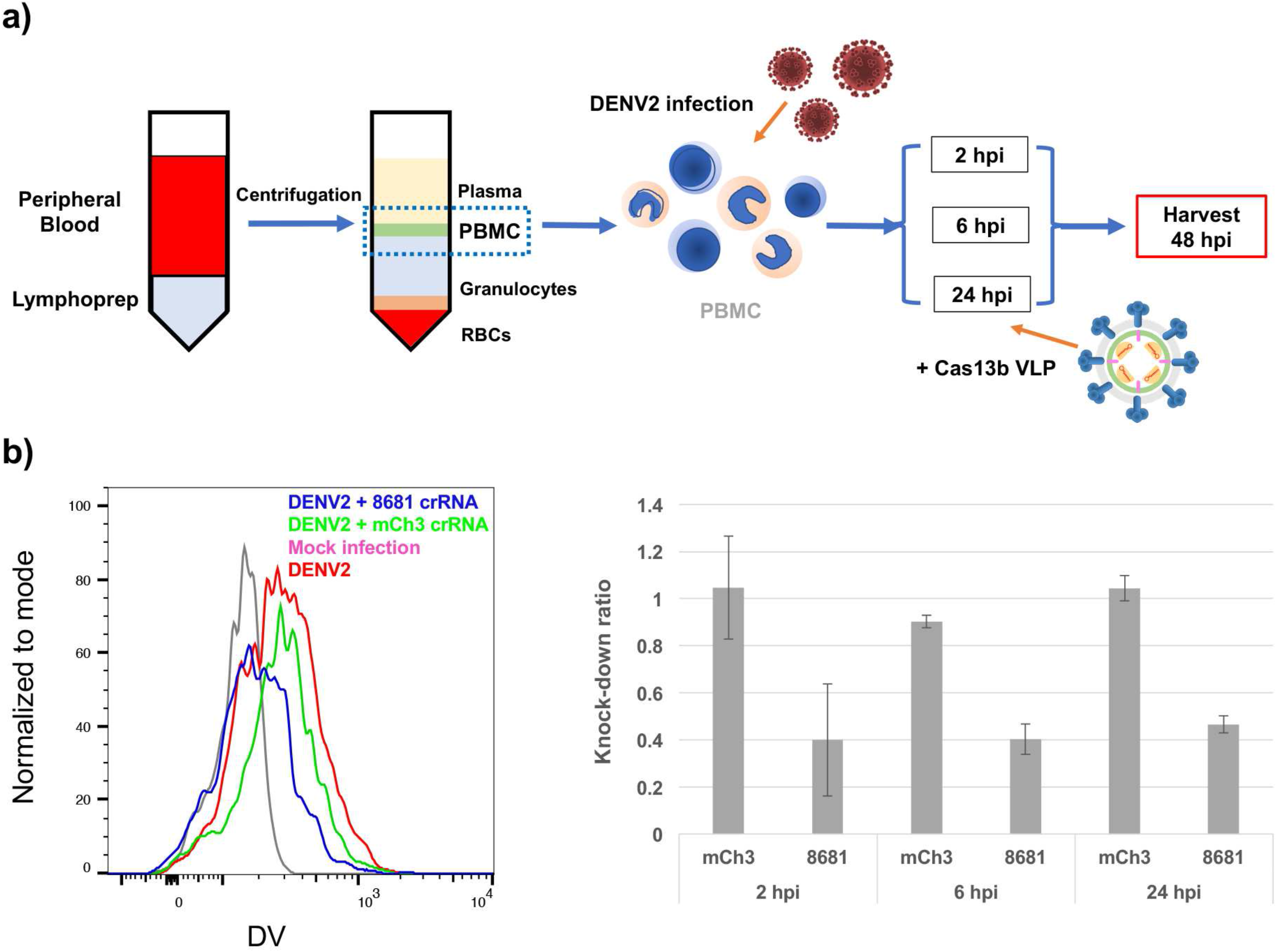
Efficient suppression of DENV2 infection in primary target cells by PspCas13b RNP delivered via virus-like particles (VLP). a) Diagram describing the experimental setup of Cas13b RNP delivery into human PBPMC infected with DENV2-16681. b) The antiviral activity of PspCas13b + 8681-nt crRNA against DENV2-16681 in PBMC. Cas13b RNP was delivered by VLP at 0, 6, or 24 hours post infection (hpi). Left histogram is a representative result from PBMC of one donor at 24 hpi VLP delivery. DV = the level of dengue E antigen measured by anti-E 4G2 mAb. Right bar plot displays DENV2 suppression with 8681 crRNA and mCh3 crRNA (nontarget control). The VLPs were delivered with equal Cas13b dose of ~77ng/100,000 cells. Knock-down ratio was calculated by the frequencies of infected cells of the VLP condition divided by the frequency of infected cells without VLP. The results represent the data from two donors.

Our results in **Supplementary Figure 1e** also suggested that the knock-down efficiency could be improved for several targets. Using BHK21-Cas13b, we tested whether spacer length could affect viral suppression efficiency for a subset of these crRNAs. Strikingly, shortening spacer length to 26-18 nucleotides of mCh3 crRNA could drastically enhance the suppression of DENV2-mCherry with maximum knock-down activity achieved between 20 and 22 nucleotides and without cross suppressing nontarget DENV2-mAmetrine (**Figure 2a**). We also found that 22-nucleotide spacer length could activate a ZIKV NS2A target region that was inactive against any crRNAs with 30-nucleotide spacer (30-nt crRNA) tiled against it (**Figure 2b**). None of the 30-nt crRNAs could suppress ZIKV beyond knock-down ratio of 0.8, while at least three 22-nt crRNAs could go beyond knock-down ratio of 0.5 (**Figure 2b**). Shortening spacer enhanced the knock-down activity of several crRNAs with varying efficiency (**Supplementary Figures 3a-b**). Nevertheless, the knock-down activity of two crRNAs (mAmet1 and 8681) were aggravated by 22-nt spacer (**Supplementary Figures 3a-b**). We selected three pairs of 30-nt vs. 22-nt crRNAs for testing by VLP delivery in BHK21 cells. The knock-down enhancement was recapitulated with Cas13b RNP delivery by VLP for 22-nt vs. 30-nt mCh3 crRNAs (**Figure 3c**). In contrast to the results in BHK21-Cas13b, the 22-nt mAmet-1 crRNA could knock down DENV2-mAmet with much less PspCas13b than its 30-nt counterpart (**Figure 3c**). The dose-response curves indicated that the 22-nt mCh3 and mAmet1 crRNAs needed 15-20 folds less of PspCas13b than 30-nt crRNAs to achieve the same level of virus suppression (**Figure 2c**). 22-nt 8681 crRNA also did not show any deterioration of knock-down activity as observed in BHK21-Cas13b (**Supplementary Figure 3b** and **Figure 2c**). These results together show that short spacer length could enhance the antiviral activity of PspCas13b and did not compromise its specificity.

**Figure 2:**
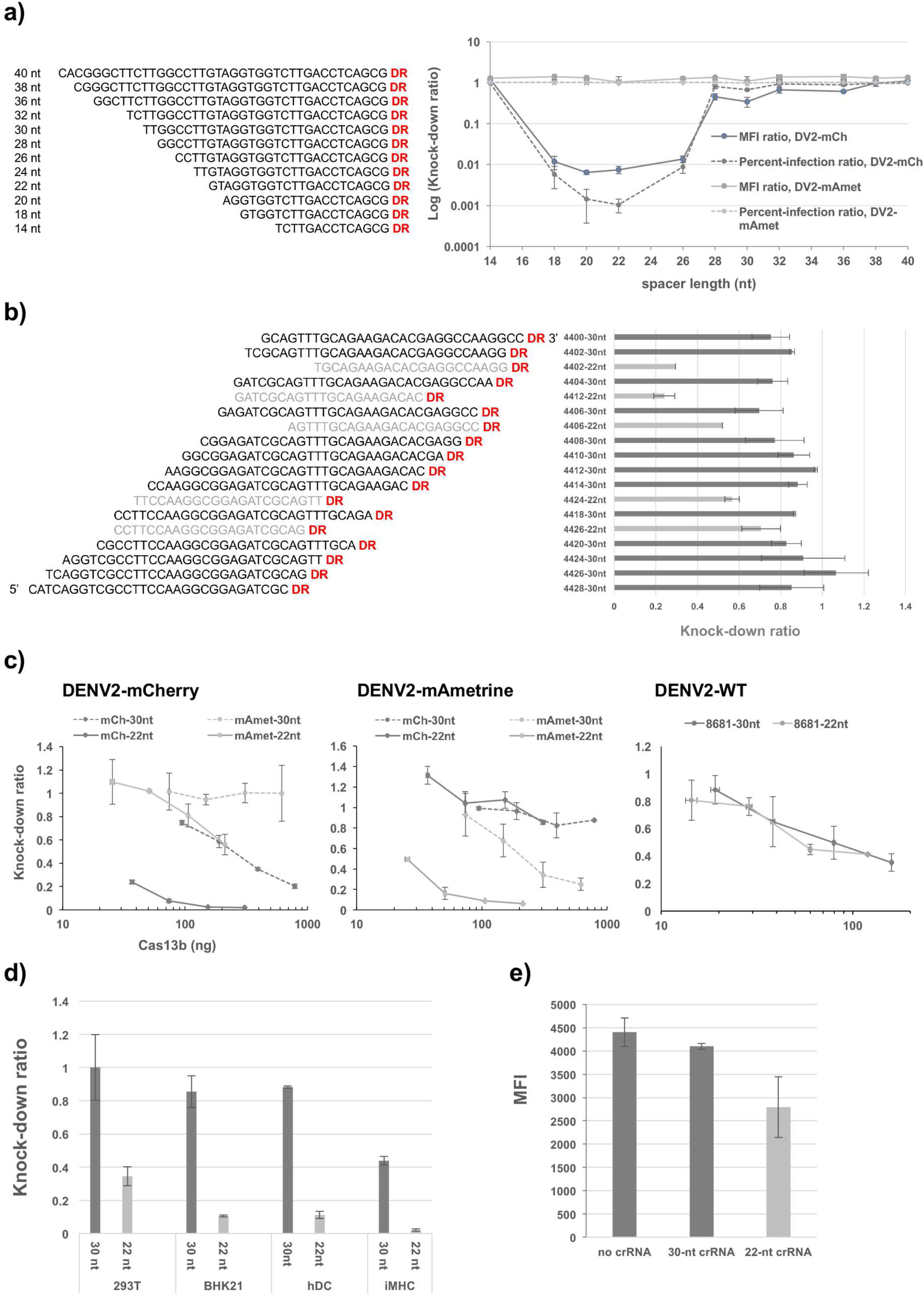
The effect of spacer length on the knock-down activity by PspCas13b. a) The effect of spacer length on antiviral activity of mCh3 crRNA against DENV2-mCherry and DENV2-mAmetrine. The right panel shows the tiled spacer sequences of mCh3 crRNAs. The right plot display the knock-down level of each DENV2 reporter with mCh3 crRNA of different spacer length. b) crRNAs with 22-nucleotide spacer could enhance the accessibility of a target region on ZIKV NS2A gene. c) Dose-response curve of Cas13b RNP delivered by VLP in BHK21 cells for three pairs of 22-nt vs. 30-nt crRNAs (mCh3, mAmet1, and 8681). d) The enhancement effect by 22-nt spacer on the antiviral activity of the mCh3 crRNA was observed in multiple cells such as 293T, BHK21, hDC, and iMHC. e) The enhancement effect by 22-nt spacer of the mCh3 crRNA on knock down of overexpressed mCherry reporter gene. The measurements were done in triplicate for a) and duplicate for b)-e). Error bar = standard deviation.

**Figure 3:**
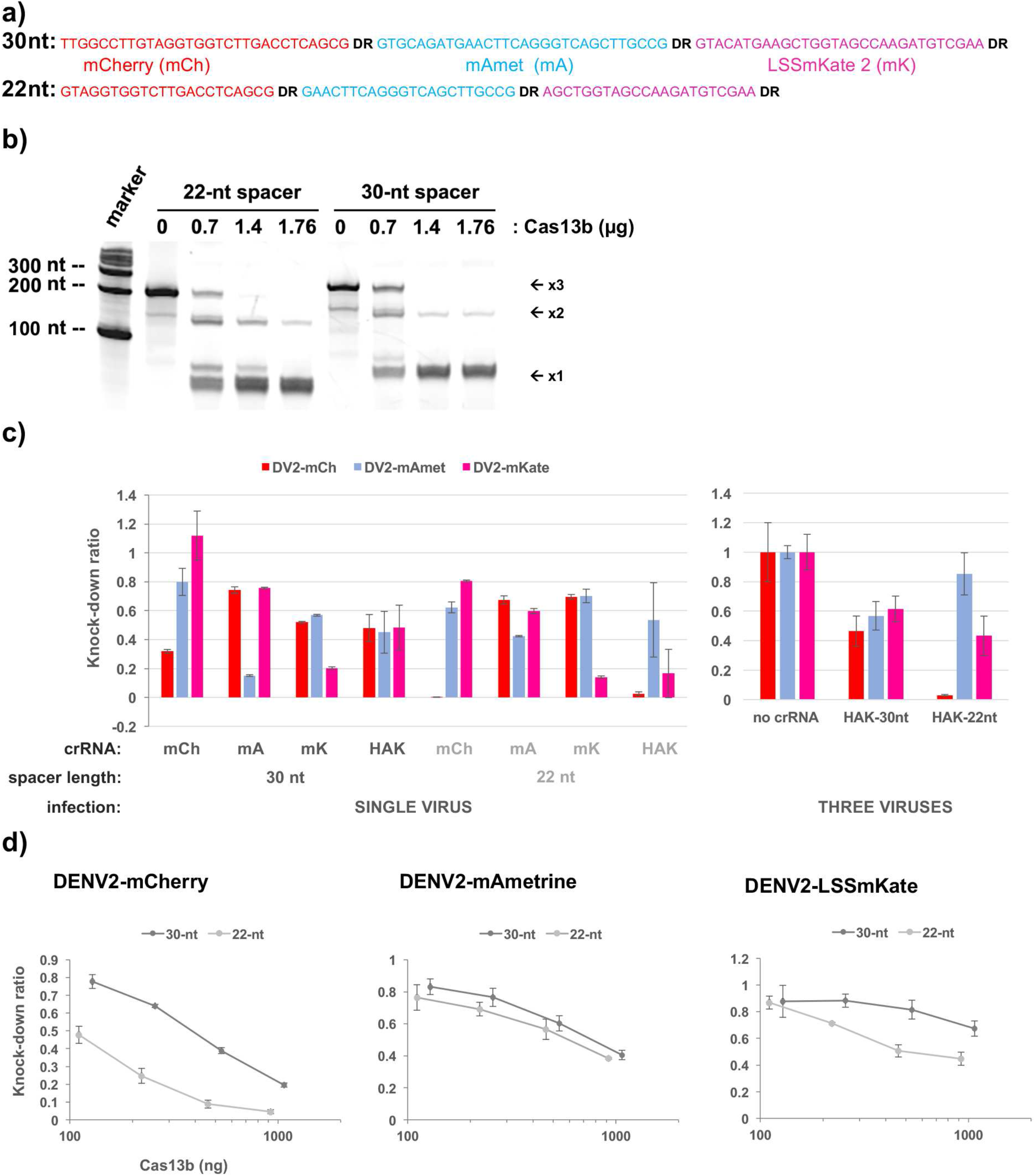
Short spacer did not inhibit crRNA processing and multiplex targeting of PspCas13b by a crRNA array. a) The sequences 30-nt and 22-nt crRNA arrays used for testing in vitro crRNA processing and multiplex targeting. DR = direct repeat. b) In vitro crRNA processing analyzed by 8M urea PAGE. 3x = crRNA array with three crRNAs, 2x = crRNA array with two crRNAs, 1x = singlet crRNAs. c) Targeting of multiple reporter DENV2 with the crRNA array in single-virus (left bar plot) and three-virus (right bar plot) infections. HAK = crRNA array shown in a). The measurements for crRNA arrays were done in triplicate. The rest were done in duplicate. d) Dose-response curves of multiplex VLP-Cas13b RNP generated with a crRNA array. The measurements were done in duplicate. Error bar = standard deviation.

We tested whether the enhancement effect of short spacer could be observed in different settings. VLP delivery of Cas13b with 22-nt mCh3 crRNA to 293T, hDC, and iMHC showed superior knock-down activity over its 30-nt counterpart with Cas13b dose at 126ng/100,000 cells (**Figure 2d**), indicating that the effect was not cell-specific. In BHK21-Cas13b, 22-nt mCh3 crRNA could knock down overexpressed mCherry reporter gene while 30-nt crRNA could not (**Figure 2e**), suggesting that the effect was applicable to regular mRNA. Interestingly, in vitro cleavage of mCherry RNAs by PspCas13b was aggravated by 22-nt crRNA (**Supplementary Figure 3c**), suggesting that the effect was specific to mammalian cells.

Since natural spacer lengths found in crRNA array of CRISPR-Cas13b in bacteria are no shorter than 30 nucleotides(Cox et al., 2017; Smargon et al., 2017), we tested whether 22-nt spacer could affect crRNA processing and multiplex targeting by a crRNA array (**Figure 3a**). In vitro crRNA processing of 22-nt and 30-nt arrays with PspCas13b were equally efficient (**Figure 3b**). Multiplex targeting of fluorescent reporter DENV2 could also be achieved with 22-nt crRNA array in both BHK21-Cas13b (**Figure 3c**) and in VLP formats (**Figure 3d**). Multiplex targeting by both 30-nt and 22-nt crRNA arrays reduced knock-down efficiency for each reporter virus slightly compared to the single-virus targeting with single crRNAs (**Figure 3c**, left bar plot). 22-nt crRNA array generally suppressed reporter DENV2 more than 30-nt crRNA array in both single-virus (**Figure 3c**, left bar plot) and triple-virus (**Figure 3c**, right bar plot) infections. Dose-response curves of multiplex VLP generated with a crRNA array showed that more Cas13b was required to achieved the same level of suppression compared to Cas13b-single crRNA by 3-10 folds (**Figure 3d** vs. **Figure 3c**). 22-nt crRNAs still suppressed viruses more than their 30-nt counterparts, though the enhancement effect of 22-nt spacer for mAmeterine was drastically reduced in the multiplex targeting (**Figure 3d** vs. **Figure 2c**).

## Discussion

In summary, we characterized the antiviral efficiency of CRISPR-Cas13b in suppressing DENV2 and ZIKV infection in mammalian cells (**Supplementary Figure 1**). We showed that transient Cas13b RNP delivery by VLP could effectively suppress dengue infection in both cell lines and primary human target cells (**Figures 1b and 2d**). We found that shorter spacer length could significantly improve the knock-down activity of CRISPR-Cas13b in mammalian cells and increased the number of targetable sites (**Figure 2**). The short spacer length (22-nt) did not affect crRNA processing and still enabled multiplex targeting with crRNA array (**Figure 3**).

Our results demonstrated the effectiveness of transient delivery of PspCas13b RNP by VLP in reducing dengue infection in human primary target cells. VLP delivery of Cas13b RNP delivery could negate several limitations such as genotoxicity and gene-size restriction imposed by gene delivery vectors such as lentivirus and AAV. Previous studies have shown that Cas13 RNP or RNA delivery could suppress RNA virus infection but the delivery methods by electroporation and transfection are not suitable for in vivo and could be highly toxic to primary cells (Bawage et al., 2018; Freije et al., 2019). Our results show that VLP delivery could effectively deliver Cas13b RNP into a variety of cells without high toxicity. Our results showed that with the right crRNAs, a picomolar range of PspCas13b could achieve strong suppression of dengue infection (e.g. 30 ng of PspCas13b per 100,000 cells (or 452 pM of PspCas13b) with 22-nt mCh3 crRNA and 22-nt mAmet1, **Figure 2c**). In case of primary human PBMC, we were able to halve DENV2 infection with ~77 ng of Cas13b/100,000 PBMC (or IC50 ~ 1.16 nM) even during ongoing infection (**Figure 1b**). The effective dose of VLP-PspCas13b RNP could rival some of the best performing small-molecule antivirals, but with added benefit of programmability.

Our results highlight a distinct difference between CRISPR-Cas13b RNA targeting in mammalian cells and in vitro/bacteria. Consistent with our study, shortening spacer length between 18-26 nucleotides did not appear to reduce PspCas13b binding to target RNA in mammalian cells as shown by previous live-imaging study using dCas13b-fluorescent fusion proteins(Yang et al., 2019). In contrast, our results of in vitro cleavage by PspCas13b (**Supplementary Figure 3c**) and previous characterizations of other Cas13b by in vitro cleavage and in bacteria showed that optimal spacer length coincides with natural spacer length of 30 nucleotides(Gootenberg et al., 2018; Smargon et al., 2017). In contrast to Cas13b and Cas13a, Cas13d naturally utilizes varying spacer lengths and was not affected by short spacer length in all settings (Konermann et al., 2018; Wessels et al., 2020; Yan et al., 2018). Despite the general benefits of short spacer lengths for PspCas13b in mammalian cells, there were cases where they aggravated knock-down activities. These exceptions were noticed only in the setting of overexpressed Cas13b and crRNA in BHK21-Cas13b (**Supplementary Figure 3**), but not in the setting of VLP delivery (**Figure 2c**). There were notable differences between the two delivery methods such as the dosage and the dynamics of Cas13b and crRNA in the cells. In VLP delivery, crRNA was pre-assembled with Cas13b before introduction into target cells while unbound crRNA could be present in the overexpression setting. In VLP delivery, the CRISPR-Cas13 components may stay only in the cytoplasm of targeted cells while crRNA in overexpression setting were produced in nucleus of targeted cells. Additional investigations will be needed to understand how these differences could affect the RNA targeting by CRISPR-Cas13b as they are relevant for designing effective delivery strategies.

Previous crRNA-scanning studies have shown that only a small fraction of target regions on mRNA in mammalian cells was targetable by CRISPR-Cas13(Abudayyeh et al., 2017; Freije et al., 2019; Wessels et al., 2020). These studies utilized a tiling crRNA library with a fixed spacer length to reveal the factors that determined the knock-down efficiency of a crRNA. Our results (**Figure 2b** and **Supplementary Figure 3**) suggest that varying spacer length in a scanning analysis could improve the number of targetable regions for CRISPR-Cas13b and might uncover additional factors that determine the knock-down efficiency of a crRNA. Combining this strategy with targeting conserved sites on viral RNA(Abbott et al., 2020; Freije et al., 2019) has the potential to identify highly potent antiviral crRNAs.

Several studies have demonstrated the use of CRISPR-Cas13 to target both plant(Aman et al., 2018a; Mahas et al., 2019) and animal(Abbott et al., 2020; Bawage et al., 2018; Freije et al., 2019) RNA viruses. Our study contributes the implementation of potent virus inhibition by PspCas13b RNP with short spacer and transient delivery by VLP in human primary cells. Our results are relevant to several emerging RNA viruses such as SARS-CoV2 that replicates in the cytoplasm. A long line of gene therapy studies to treat cystic fibrosis using lentivirus delivery in vivo supports the possibility of using pseudotyped Cas13b-VLP to treat viral infection in lung and respiratory track(Marquez Loza et al., 2019). In addition to RNA viruses, our results are applicable to knocking down specific host genes in primary cells for transcriptome engineering.

## Materials and Methods

### Cells and culture media

BHK21, BHK21-rtTA3(Suphatrakul et al., 2018), and 293T were cultured in high-glucose DMEM (HyClone) supplemented with 10% heat-inactivated fetal bovine serum (HI-FBS, ThermoFisher Scientific), 100 U/ml penicillin-G, and 100 μg/ml streptomycin sulfate (ThermoFisher Scientific) (D10). Vero cells was maintained in MEM (ThermoFisher Scientific) supplemented with 10% HI-FBS and 100 U/ml penicillin-G, and 100 μg/ml streptomycin sulfate (MEM10). iMHC was maintained in DMEM-F12 (Hyclone) supplemented with 10% HI-FBS). C6/36 was maintained in Leibovitz L-15 (HyClone) supplemented with 10% HI-FBS, 10% tryptose phosphate broth (Sigma), and 100 U/ml penicillin-G, and 100 μg/ml streptomycin sulfate. Human primary PBMC, macrophages, and CD14^+^ monocytes were maintained in RPMI-1640 (HyClone) supplemented with 10% HI-FBS, 100 U/ml penicillin-G, and 100 μg/ml streptomycin sulfate (R10). All the adherent cells in this study were detached by 0.1% Trypsin-EDTA (ThermoFisher Scientific). All mammalian cells were cultured in 5% CO_2_ at 37°C with at least 80% humidity. C6/36 was cultured at 28°C.

Primary human PBMC were obtained from fresh blood of healthy volunteers. Human blood was obtained from donors after providing informed consent, following a protocol (Siriraj-IRB COA no. Si707/2016, Protocol number: 632/2559 (EC2)) approved by Faculty of Medicine, Siriraj Hospital, Mahidol University, Thailand. The blood was diluted 1:1 ratio of normal saline and layered over Lymphoprep™ (Axis-shield) gradient density at 1,077 g/ml. After centrifugation at 800 x g for 25 minutes (min) at room temperature (RT), plasma samples were removed from the top of the solution and PBMC were recovered from the underlying layer. The PBMC were washed twice in RPMI 1640 medium and then incubated with red blood cells lysis buffer at room temperature for 5 minutes to lyse contaminating RBCs. The PBMC were washed once in R10. PBMC viability, measured by trypan blue exclusion, was greater than 95%.

Human dendritic cells (hDC) were derived from PBMC according to published protocol (Boonnak et al., 2011). 1.5 × 10^7^ of PBMC were seeded on each 60 mm-Primaria plate (Corning) and cultured for 1 hour. Cells were washed with 5 ml of plain RPMI, 10 times to remove non-adherent cells. Washed PBMC was then replenished with 5 ml of R10 supplementing with 2,000 U/ml of rh GM-CSF (ThermoFisher Scientific) and 4,000 U/ml of rh IL-4 (R&D Systems) and cultured for 5 days. DC were then harvested from culture supernatant for experiments and analysis. DC were verified with a panel of CD14-PerCP (Miltenyl Biotech), CD83-BV510 (BD Pharmingen), CD86-PE (BD Pharmingen), HLA-DR-APC-Vio770 (Miltenyl Biotech), CD209-PE-Vio770 (Miltenyl Biotech), and CD163-FITC (BioLegend) mAbs by flow cytometry (BD LSRFortessa).

Fresh PBMC were used to isolate primary monocytes using Dynabeads® Untouched™ Human Monocytes (ThermoFisher Scientific) according to the manufacturer’s instructions. Briefly, the PBMC were incubated with blocking reagent and antibody mixture for 20 min at 2 to 8 °C to label cells which were not monocytes. The labeled cells were mixed with Dynabeads® and incubated for 15 min at 2 to 8°C. To remove non – monocytes, the labeled cells were placed in a magnet (DynaMag™). The purity of the isolated monocyte subsets, evaluated by flow cytometry with CD14-PerCP mAb (Miltenyl Biotech), was consistently 90 to 95 % of CD14^+^ cells (Boonnak et al., 2011).

The isolated CD14^+^ monocytes were cultured in R10 supplemented with 5% HI-human AB serum to allow differentiation into macrophages(Brugger et al., 1991). Half volume of medium was replaced with fresh medium every 3 days. The macrophages were harvested for experiments on day 13. The macrophages were then verified by flow cytometry analysis with CD163-FITC mAb (Biolegend).

### Plasmids

Spacer sequences used to construct crRNAs are listed in **Supplementary table 1**, **Figure 2**, and **Supplementary Figure 3**. All the plasmids were constructed by either Gibson assembly or standard T4 ligation.

PspCas13b from pC0046(Cox et al., 2017) (Addgene #103862) was subcloned to replace CasRx on pXR001(Konermann et al., 2018) (Addgene #109049). The pXR001-PspCas13b was then engineered to replace HIV NES with VAMP2 (amino-acid sequence = KTGKNLKMMIILGVICAIILIIIIVYFTGSR) or SQS (amino-acid sequence = SRSHYSPIYLSFVMLLAALSWQYLTTLSQVTED) to generate lentiviral plasmids for PspCas19-VAMP2 and PspCas13-SQS. These pXR001-Cas13b plasmids were used for the experiment in Figure 1b.

To create lentiviral vector with inducible expression of Cas13b, PspCas13b from pC0046(Cox et al., 2017) (Addgene #103862) was subcloned into pENTR1A(Campeau et al., 2009) (Addgene #17398) at EcoRI site. PspCas13b on pENTR1A was shuttled to pLenti-CMVtight-Blast-DEST(Campeau et al., 2009) (Addgene #26434) with Gateway cloning (ThermoFisher Scientific). Blasticidin resistant gene was then replaced with eGFP for the purpose of cell sorting. This pLenti-PspCas13b plasmid was used to generate lentivirus for deriving BHK21-Cas13b stable cell line.

pBA439(Adamson et al., 2016) (Addgene #85967) was engineered to replace Cas9-gRNA cassette with Psp direct repeat (DR) with BsmBI sites upstream for cloning spacer and to replace BFP with miRFP703 for the purpose of cell sorting. The lentiviral plasmid, pBA439-Psp-miRFP was then used for testing crRNAs in BHK21-Cas13b.

To create plasmid for PspCas13b RNP delivery by VLP, BIC-Gag-CAS9(Mangeot et al., 2019) (Addgene #119942) was engineered to replace Cas9 with PspCas13b-HA to generate BIC-Gag-PspCas13b. pC0043 (Addgene #103854) was used to clone spacer to generate a crRNA for VLP production.

PspCas13b on pC0068 plasmid (Addgene # 115219) was replaced with PspCas13b with HIV NES and HA tag (from pC0046 (Addgene #103862)) to generate pC0068-PspCas13b-HA for PspCas13b-HA purification.

crRNA arrays (mCh3-mAmet1-mK1) was generated from T4 ligation of 3 pairs of annealed primers with asymmetric, unique overhangs to pBA439-Psp-miRFP linearized with BsmBI for experiments in BHK21-Cas13b. The crRNA arrays were then subcloned onto pC0043 linearized with BbsI and XhoI to generate pC0043-crRNA array plasmid for multiplex VLP generation.

### Viruses

DENV2-16681 and ZIKV-SV0010/15(Buathong et al., 2015) were produced from C6/36 cell line cultured in L-15 (HyClone) supplemented with 1.5% HI-FBS (ThermoFisher Scientific), 10% tryptose phosphate, and 100 U/ml penicillin-G, and 100 μg/ml streptomycin sulfate. The infectious titers were quantitated by foci assay in Vero cells stained with anti-E 4G2 monoclonal antibody as previously described(Suphatrakul et al., 2018). Fluorescent reporter DENV2 were generated and quantitated as previously described(Suphatrakul et al., 2018).

### Lentivirus for gene delivery

Pseudotyped lentivirus particles were generated by co-transfection of lentiviral plasmids (e.g. pBA439-Psp-miRFP and pLenti-CMVtight-PspCas13b-eGFP) with pCMV-VSV-G(STEWART, 2003) (Addgene #8454) and psPAX2 (Addgene #12260) using PEI into 293T cells cultured in D10. The media was harvested for lentivirus particles 2 days post transfection.

### Immunofluorescent microscopy

Briefly, cells were seeded in a 96 well plate to reach 50-60 % confluency overnight. Cells were washed and fixed with 100 μL 4% paraformaldehye in PBS for 10 minutes at 37°C, permeabilized with 100 μL 1% Triton X-100 in 1x PBS for 15 minutes at 37°C, and blocked with 1% BSA in 1x PBS for 1 hour at 37°C. Cells were then incubated with primary antibody (HA-Tag Mouse mAb Cell Signaling, 1:100 dilution), washed and incubated with secondary antibody (Cy3 conjugated goat anti-mouse (Jackson ImmunoResearch), 1:2000 dilution). Samples were washed and images were obtained using EVOS fluorescence microscope (ThermoFisher Scientific).

### Testing crRNAs with Cas13b inducible expression system in BHK21

BHK21-rtTA3 cell line(Suphatrakul et al., 2018) was transduced with lentivirus generated from pLenti-PspCas13b. Transduced cells (GFP^+^) were sorted into single cells to isolate stable clones on BD FACSAria III. To screen for desired clone (BHK21-Cas13b), we tested the clones for uniform Cas13b expression and their abilities to knock down DENV2-mCherry with mCh3 30-nt crRNA. A BHK21-Cas13b clone was selected for all subsequent experiments. crRNA was introduced into the BHK21-Cas13b clone with the lentivirus carrying mU6-crRNA expression cassette and miRFP703-2A-puromycin cassette for cell sorting. Cells that were positive for both GFP (PspCas13b) and miRFP703 (crRNA) were sorted as a pool. The pool cells were maintained for at least a week under D10 + 5 μg/ml puromycin before the knock-down experiments. To test for knock-down, 100,000 cells/well were seeded in 6-well plate with D10 + 0.1 μg/ml of doxycycline to induce Cas13b expression. After 24 hours, the induced cells were then infected with DENV2 or ZIKV at M.O.I. = 0.1 to test for knock-down activity of the crRNA. The cells were maintained at 37°C for 72 days before harvested by trypsin digestion and fixed with 3.7% formaldehyde in 1x PBS for measurement of virus infection by flow cytometry (BD LSRFortessa).

Virus infection was measured by mean fluorescent intensity (MFI) of fluorescent reporters (for reporter DENV2 such as DENV2-mCherry, DENV2-mAmetrine, and DENV2-LSSmKate2) or viral E antigens (stained with 4G2 mAb for DENV2-16681 and ZIKV-SV0010/15) of total cells or percent infection of total cells. For knock-down ratio (KD ratio), MFI (or percent infection of total cells) of BHK21-Cas13b with an experimental crRNA was divided by the MFI (or percent infection of total cells) of BHK21-Cas13b with a non-targeting crRNA.

### Production of VLP-YFP and VLP-Cas13b RNP

17-million 293T cells were seeded in 150-mm dish. To generate VLP-Cas13b RNP, one dish of 293T was transfected with four plasmids 5.1 μg BIC-Gag-PspCas13b, 13.2 μg pC0043-crRNA, 3.3 μg pCMV-VSV-G (STEWART, 2003) (Addgene #8454) and 8.4 μg pBS-CMV-gagpol (Addgene #35614) using 90 μg 1mg/ml PEI (3ug PEI: 1ug total DNA) 24 hours after seeding. For VLP-YFP, BIC-Gag-PspCas13b and pC0043-crRNA were replaced with 5.1 μg MLV-Gag-YFP(Sherer et al., 2003) (Addgene #1813). The transfected cells were maintained for 2 days before harvest of media. Harvested media was clarified by centrifugation at 1000 x g, 4°C, 10 minutes and then filtered with 0.45-μm PES-membrane syringe filter (Millipore). 30 ml of clarified media was then layered on 10 ml of 10% w/v sucrose cushion in 1xPBS in 50-ml falcon tube (Corning) and centrifuged at 10000 xg, 4°C for 4 hours in JA14 rotor. The supernatant was carefully discarded and dried by pressing against paper towel for 30 seconds. The pellet was resuspended with D10. Dissolved pellet was aliquoted and stored frozen at −70°C.

### Purification of PspCas13b and quantitation of PspCas13b in VLP

PspCas13b-HA was expressed and purified as previously reported with some modifications(Gootenberg et al., 2018). PspCas13b-HA was expressed in BL21-Rossetta2 by 2mM IPTG induction in 2xYT media at O.D. of 0.6, 22°C for 18 hours. The bacteria cells were harvested by centrifugation and stored at −70°C until purification. The bacteria pellet was resuspended in 20mM Tris-HCl pH 8.0, 500mM NaCl, 1mM DTT + protease inhibitors (EDTA-free, Roche) + lysozyme, lysed by sonication, and clarified by centrifugation at 10,000 xg, 4°C for 20 minuted. Clear lysate was loaded onto streptactin column. The column was then washed with 20mM Tris-HCl pH 8.0, 500mM NaCl, 1mM DTT before elution with the same buffer + 0.15% NP-40 + 2.5 mM desthiobiotin. The eluted fraction was treated with Ulp1 protease to cleave affinity tag overnight at 4°C. Ulp1 and cleaved affinity tag were removed by NiNTA (Roche). The NiNTA flow through was adjusted to 250mM NaCl and purified on HiTRAP-SP (20mM HEPES pH 7.3, 250mM NaCl, 5% glycerol, 1mM DTT). We obtain highly purified PspCas13b-HA that was used as Cas13b reference for VLP quantitation and in vitro crRNA processing.

To quantitate for Cas13b in VLP, 2 μl from serial dilutions of pure PspCas13b-HA and VLP were dotted on a strip of nitrocellulose membrane. The membrane was blocked with 5% skim milk in 1xPBS-T. The blocked membrane was then probed with anti-HA (1:1000 dilution, CellSignalling) and rabbit anti-mouse P260 IgG conjugated with HRP (1:1000 dilution, DAKO). The antibody-stained membrane was then soaked with SuperSignal West Pico PLUS substrate (ThermoFisher Scientific) according to manufacturer’s instruction and imaged on C-Digit blot scanner (LI-COR). Quantitation of signal was performed with Image Studio Lite program (LI-COR).

### Delivery of protein cargo by VLP

VLP delivery media was prepared by mixing a volume of VLP preparation according to the desired Cas13b dose and topped up with D10 to 100 μl. For VLP-YFP, 50 μl of VLP preparation was mixed with 50 μl D10. To deliver VLP into BHK21, iMHC, and 293T cells, the cells (BHK21 ~100,000 cells; 293T ~150,000-200,000 cells) in one well of 24-well plate were treated with 300 μl of complete media + 8 μg/ml polybrene for 10 minutes at 37°C before adding 100 μl of VLP delivery media. For these cells, the VLP-treated cells were then maintained for 24 hours at 37°C before exchanging media to 500 μl D10 for another 24 hours before harvest for analysis. To deliver VLP into hDC, 50,000 cells of hDC were centrifuged at 450xg, 5 minutes, 4°C and the cell pellet was resuspended with 300 μl R10+ 8 μg/ml polybrene. 100 μl of VLP delivery media (R10) was added to hDC. The cells were then transferred to one well of 24-well plate and cultured for 48 hours at 37°C before harvest. VLP delivery of YFP was performed with the same protocol as VLP delivery of Cas13b.

### Test of knock down in hDC, 293T, iMHC by VLP

For iMHC and BHK21, 25,000 cells/well were seeded on 24-well plate. For 293T, 50,000 cells/well were seeded on 24-well plate. After 24 hours, the cells were infected with DENV2-mCherry at MOI 0.1 for another 24 hours. VLP was then delivered as detailed above. For hDC, cell suspension (~50,000 cells/20 μl) was mixed 30 μl of DENV2-mCherry (MOI = 1.0) and topped up with 50 μl R10 in 1.5-ml Eppendorf tube and incubated at 37°C for 2 hours. Then, the cells were washed once with 50 μl R10 by centrifugation at 450xg, 5minutes, 4°C and proceeded to VLP delivery.

### Test of knock down in BHK21 with overexpressed mCherry reporter

BHK21-Cas13b cell line was transduced with lentivirus carrying TRE-mCherry reporter cassette (generated from pLV-tetO-mCherry, Addgene #70273). The transduced cells were sorted for mCherry+ cells. The BHK21-Cas13b-mCherry was then transduced with crRNA lentivirus and sorted for GFP^+^mCherry^+^miRFP703^+^ cells. The sorted cells were maintained in D10+5 μg/ml puromycin for 1 week before knock-down experiment. The knock down was initiated by adding 0.1 μg/ml doxycycline and continued the culture for 48 hours. The cells were harvested by trypsin digestion and fixed with 3.7% formaldehyde in 1xPBS for analysis by flow cytometry.

### DENV2 knock down in infected human peripheral blood monocytes

PBMCs were obtained as detailed above. The experiments were performed in 24-well plate with PBMC seeded at 100,000 cells/well in R10. To infect PBMC with DENV2-16681, 0.1 mg/ml of purified 4G2 (*Flavivirus* E protein specific mAbs) were incubated with virus for one hour at 37°C before adding to PBMCs at MOI of 1 in R10. PBMC were infected for 2 hours at 37°C. To deliver VLP, 200 μl of culture media was removed and 100 μl of R10 with 8 μg/ml polybrene was added to the well. PBMC were treated with polybrene at 37°C for 10 minutes before 100 μl of VLP delivery media was added. Infected PBMC were cultured for 48 hours post infection before harvest for analysis. The culture media was also collected and clarified by centrifugation at 4°C for foci assay to quantitate infectious virus titer.

Harvested PBMCs were washed and stained with live dead dye (Invitrogen) according to manufacturer’s instruction. Stained cells were fixed and permeabilized with 3.7% formaldehyde and 0.5% saponin, respectively. Cells were then intracellularly stained with 4G2 mAbs followed by rabbit anti-mouse Igs FITC (Dako). The cells were fixed with 1% formaldehyde and analyzed on BD LSRFortessa. Data analysis was performed using FlowJo software version 10.1.

For testing the uptake of VLPs in PBMCs, cells were incubated with YFP-VLPs 50 μl for 48 hrs at 37°C. Cells were washed and surface stained with CD3 APC (BD Pharmingen), CD14 PerCP (BD Pharmingen) and CD19 PE (Dako) mAbs for 30 min. Thereafter, cells were washed, fixed with 1% formaldehyde and analyzed on BD LSRFortessa. Data analysis was performed using FlowJo software version 10.1.

### In vitro transcription and crRNA processing assay

DNA template for in vitro transcription of a crRNA array was amplified from pBA439-HAK by PCR using a forward primer with T7 promoter sequence and a reverse primer that included poly-T to terminate transcription. 0.5 μg of DNA template were transcribed in 100 μl reaction using 30 μg T7 RNAP, 2mM NTPs, 40 U RNaseIN (Promega) in 50mM Tris-HCl pH 7.5, 15mM MgCl_2_, 5mM DTT, and 2mM spermidine at 37°C for 2 hours. The transcription reaction was treated with 2 U of DNase I (ThermoFisher Scientific) at 37°C for 30 minutes and clean-up using Qiagen RNA easy kit according to manufacturer’s protocol. In vitro crRNA array processing was carried out in 10 mM Tris-HCl pH 7.5, 50 mM NaCl, 0.5 mM MgCl_2_, 20U RNaseIN (Promega), 0.1% BSA for 30 minutes at 37°C, stopped by adding 1% SDS, 2x TBE-Urea gel loading buffer and denatured for 10 minutes at 95°C. Samples were then put on ice for 10 minutes before running them on an 12% TBE-8 M Urea polyacrylamide gel in 1x TBE buffer at 200 V for 40 minutes. Gel staining was carried out in 1x Sybr Gold in 1x TBE for 5 minutes and imaged on a gel doc system.

### In vitro cleavage of target RNA by PspCas13b

Cas13b-cRNARNP was formed by mixing 1 or 2 uM of crRNA with 1.8 uM of Cas13b and incubate on ice 10 min in 5 μl water. The preformed RNP was then added to 10 μl in vitro cleavage reaction that contained 20 ng of mCherry RNA, 1 ul of 10x Cut smart buffer (NEB), 4U of RNase IN and incubated at 37°C for 2 hours. The reaction was stop with 1%SDS and then extracted by phenol/chloroform for ethanol precipitation. The RNA pellet was resuspended with 1X RNA loading buffer (4 mM EDTA, pH 8.0, 2.7% formaldehyde, 20% glycerol, 7.7 M formamide, 80 mM MOPS, 20 mM sodium acetate 0.025% (v/v) bromophenol blue) and heated at 95°C for 10 minutes and then put on ice 2 minutes before loading on 8% TBE-polyacrylamide gel with 8 M urea.

## Acknowledgement and conflict of interest

We would like to acknowledge Nuntaya Punyadee for technical assistance with human macrophages and iMHC cultures; Korbporn Boonnak for her technical advice on human dendritic cells; Sutha Sangiambut for assistance with chemiluminescent measurement. This study was funded by a research grant from Platform Technology Program, BIOTEC (P-18-52192). S.O., E.S., P.P., A.S., and B.S. are co-inventors on a patent application based on this work.

## Supplementary Figure Legend

**Supplementary Figure 1:**
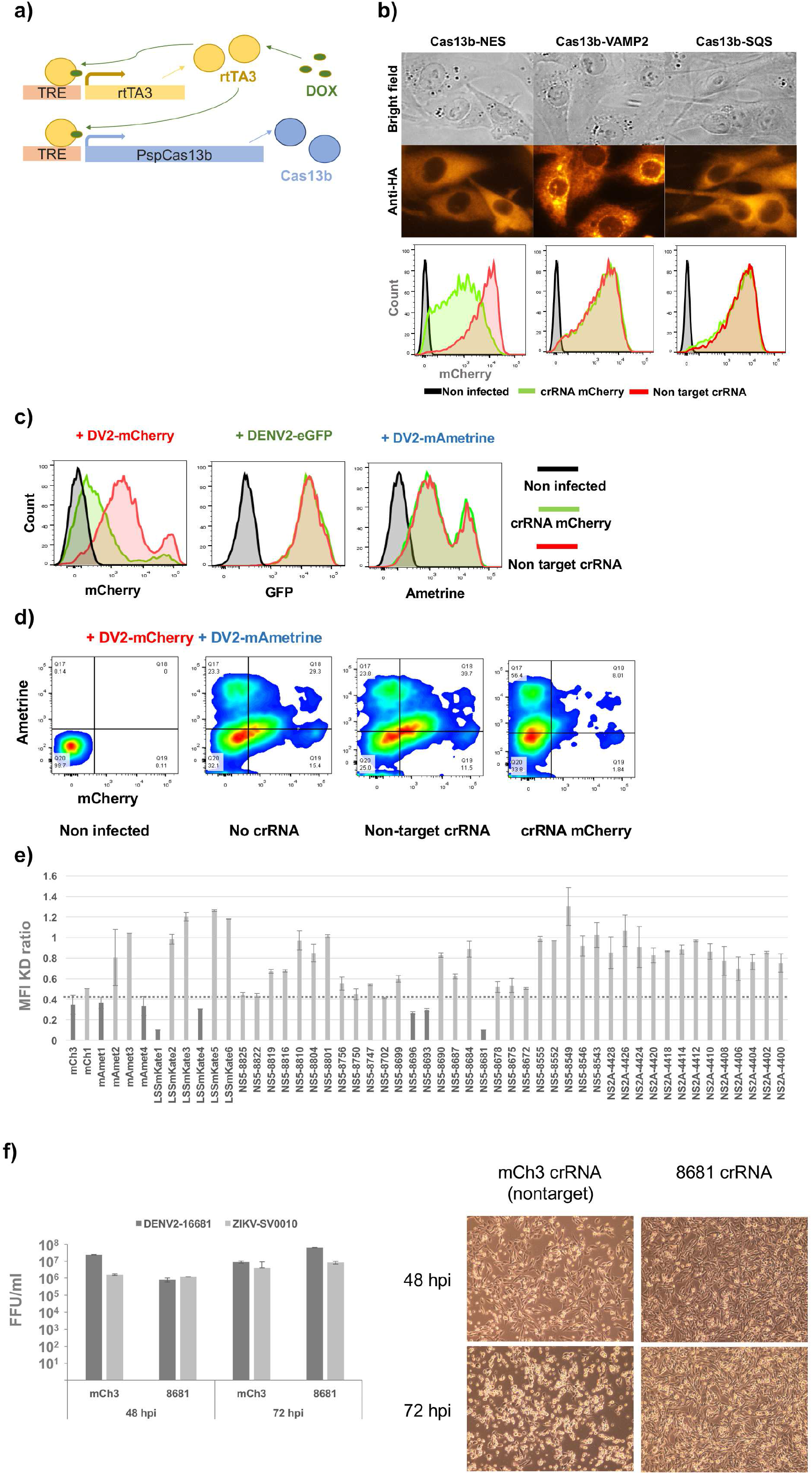
Flaviviral suppression with CRISPR-Cas13b by inducible expression in BHK21. a) Diagram describing the inducible expression system that controlled the expression of PspCas13b in BHK21 cells. b) Cytoplasmic PspCas13b, but not ER-localized PspCas13b, could suppress DENV2-mCherry replication. The top panel is the bright-field image of BHK21 expressing various form of PspCas13b. The middle panel is the same views of the top panel but with anti-HA signal representing the location of PspCas13b. The bottom panel is histograms of DENV2-mCherry intensity in BHK21 with crRNAs and different forms of PspCas13b. c) Specific viral suppression of reporter DENV2 by an mCherry-targeting crRNA (mCh3 crRNA) in single-virus infection setting. Viral suppression for each reporter DENV2 is shown as mean-fluorescent intensity (MFI) histograms from flow-cytometry measurement of infected cells. d) Specific viral suppression of reporter DENV2 by mCh3 crRNA in co-infection setting. Viral suppression for each reporter DENV2 is shown as heat-map scatter plot from flow-cytometry measurement of infected cells. e) Summary bar plot of viral suppression activities of 51 crRNAs individually tested. Viral suppression is presented as a mean of knock-down ratio (KD ratio) calculated from the ratio of MFI in BHK21-Cas13b with experimental crRNA relative to the MFI in BHK21-Cas13b with nontarget crRNA from duplicate measurements (error bar = standard deviation). Dark grey bars highlight the crRNA with KD ratio below 0.4. The details of the tested viruses and crRNA spacer sequences are listed in **Supplementary table 1**. f) Suppression of wild-type DENV2-16681 by a crRNA targeting DENV2 NS5 gene (8681 crRNA). The left bar plot compare infectious titers of DENV2-16681 between BHK21-Cas13b with 8681 crRNA and BHK21-Cas13b with mCh3 crRNA (nontarget control) at 48 hours post infection (hpi) and 72 hpi (error bar = standard deviation). The measurements were done in duplicate. The right panel is representative bright-field images of the infected cells at 48 and 72 hpi.

**Supplementary Figure 2:**
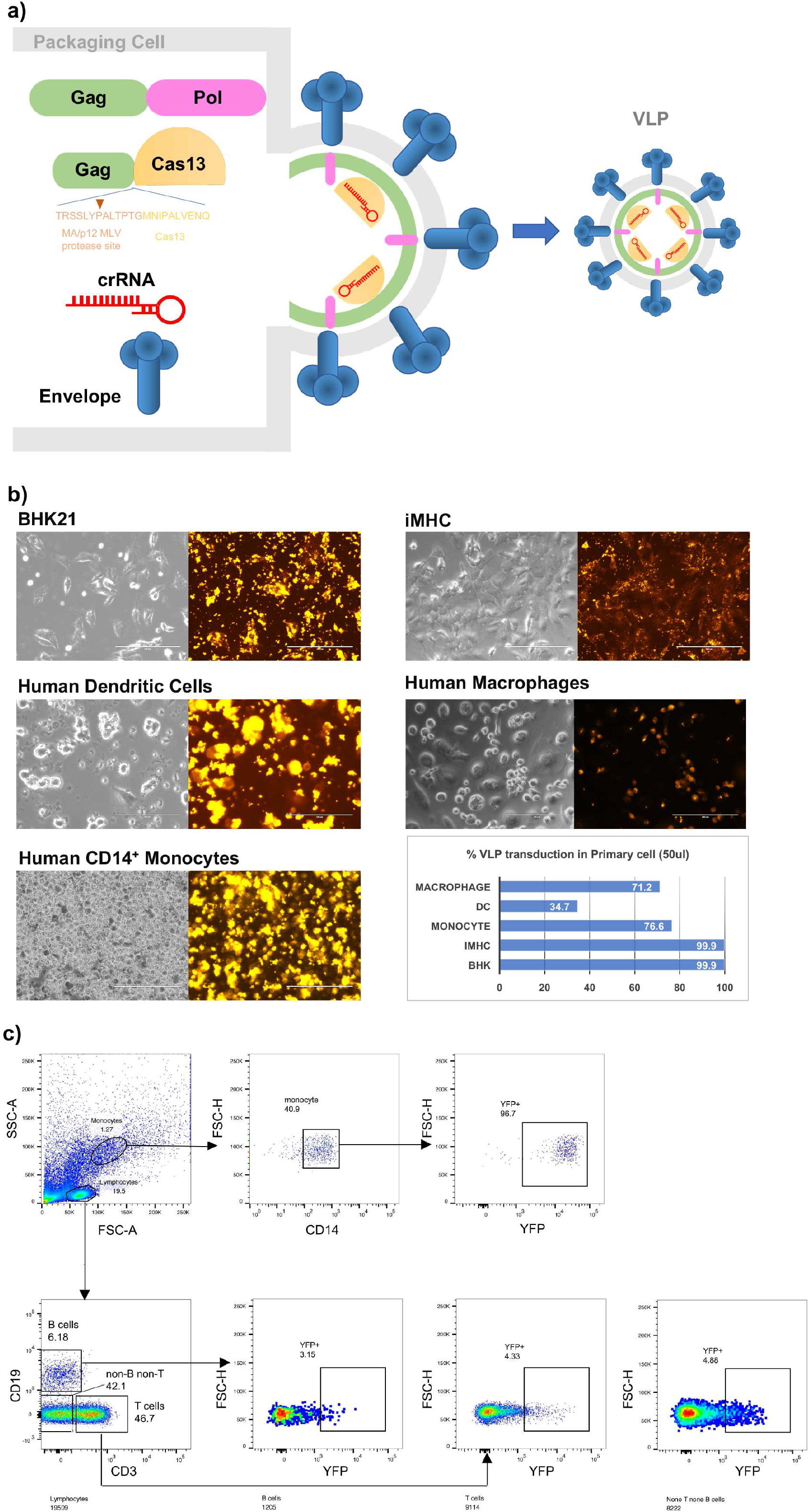
VLP delivery of protein cargo into human primary target cells. a) Diagram showing the generation of VLP for PspCas13b RNP delivery (adapted from (Mangeot et al., 2019)). b) Delivery of YFP into various mammalian cells. The figure show the results of transducing BHK21, hDC, macrophages, iMHC, and CD14+ monocytes with VLP-YFP from fluorescent microscopy (bright-field images and fluorescent images) and percent transduction measured by flow cytometry. c) Flow cytometry analysis of YFP delivery by VLP into different cell populations of human PBMC.

**Supplementary Figure 3:**
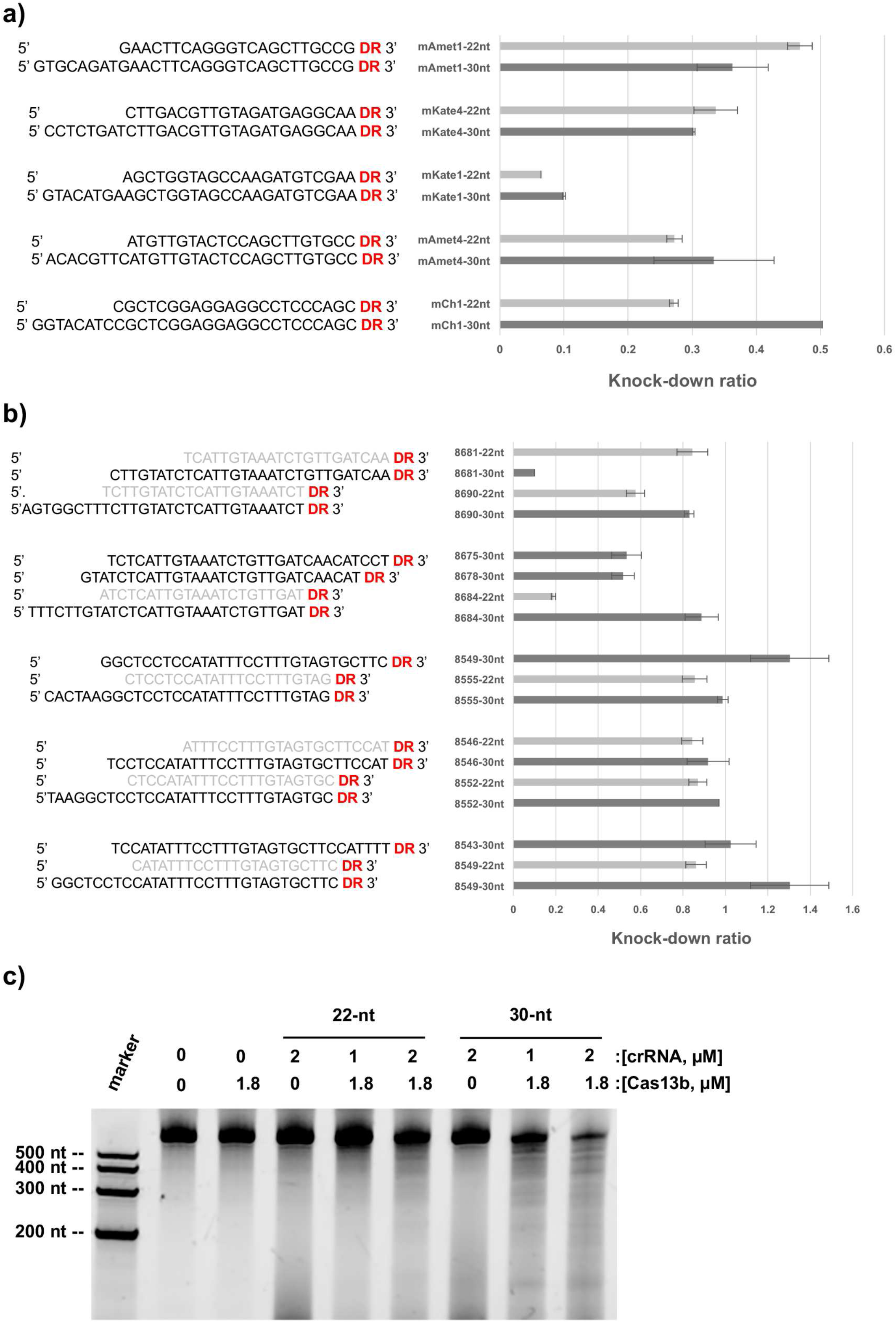
The comparison of virus suppression activity between 22-nt and 30-nt crRNAs and in vitro cleavage. a) crRNA set for targeting fluorescent reporter genes. b) crRNA set for targeting NS5 gene of DENV2-16681. **c)** In vitro cleavage of mCherry RNA by PspCas13b + 22-nt mCh3 crRNA vs. 30-nt mCh3 crRNA.

**Supplementary Table 1.**
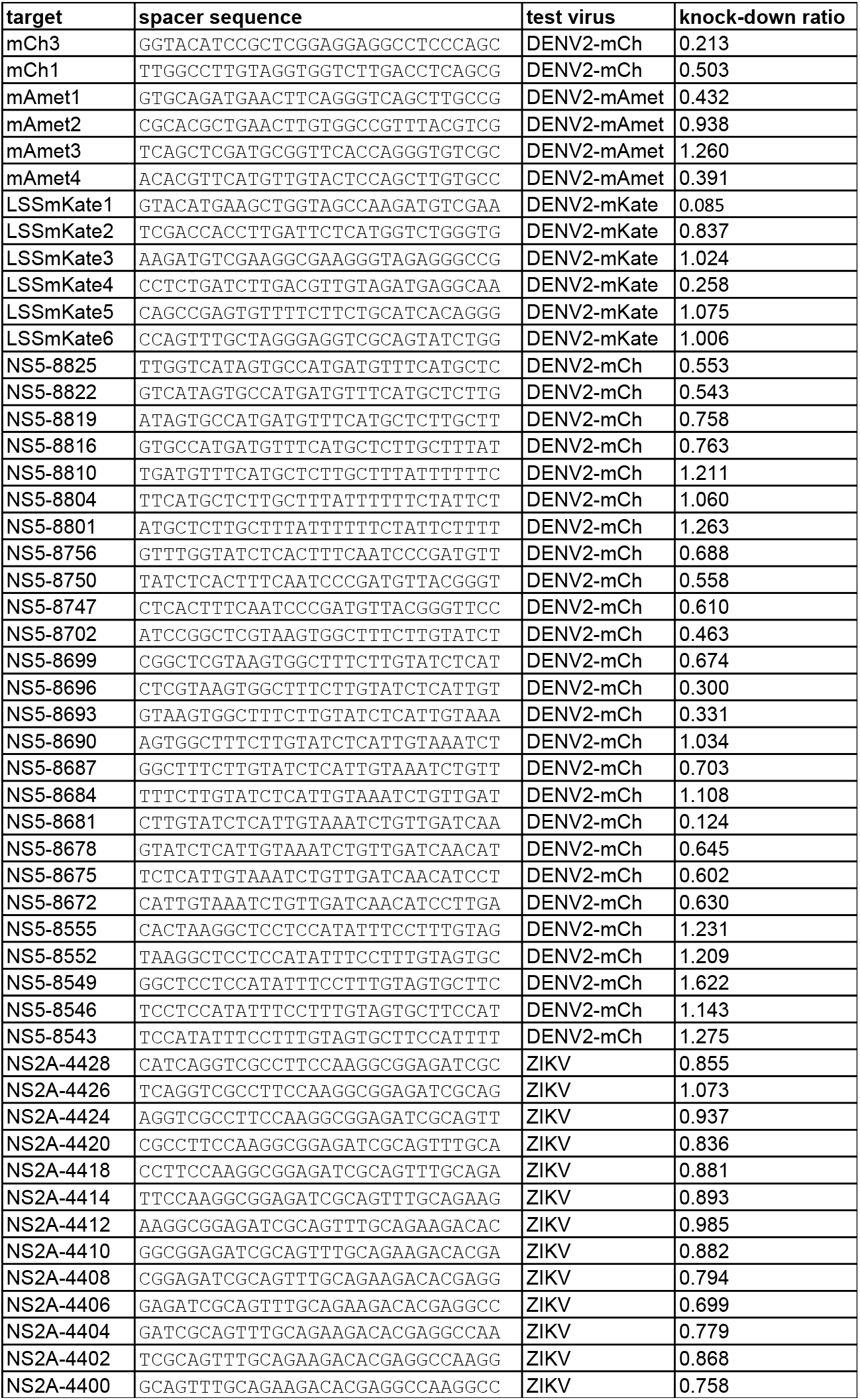

